# Effects of shade-coffee certification programs on bird, trees and butterfly diversity in Colombia

**DOI:** 10.1101/251231

**Authors:** Juan Pablo Gomez, Elena Ortiz-Acevedo, Jorge E. Botero

## Abstract

In the last decade coffee certification programs have grown rapidly in Latin America, encouraging producers to harvest coffee based on production standards intended to enhance biodiversity conservation. However, few studies have tested whether such programs have a positive conservation impact. To date, research has focused on comparing community similarity between forests and plantations, but the question of whether certified plantations provide refuges for biodiversity in regions where all the forest has been lost remains untested. Here, we compare bird, butterfly and plant communities in highly deforested regions in Santander, Colombia, to determine the potential conservation role of two certification programs: Rainforest Alliance and Rainforest Alliance+Organic. We used 13 farms to census birds, butterflies, and trees, and quantified structural characteristics of the shade. We found little difference in most measures of diversity and composition of birds, butterfly and plant communities between types of plantations. However, despite high variation across farms, butterfly richness and abundance increased with the decrease in the use of pesticides in plantations. These results suggest that reduced use of chemical compounds in certified coffee plantations might enhance conservation of butterfly communities. The biodiversity associated with these coffee plantations and the high deforestation rates in Santander, suggest that irrespective of their certification type they provide the last refuges for biodiversity conservation in this region.

## Resumen

En la ultima década, los programas de certificación al café en América latina han crecido rápidamente, motivando a los productores a sembrar café basado en estándares de producción que promueven la conservación de la biodiversidad. Sin embargo, pocos estudios han intentado comprobar si estos programas de certificación tienen un impacto positivo sobre la conservación de la biodiversidad. Hasta la fecha, la mayoría de estudios se han enfocado en comparar la similitud entre las comunidades de bosques y plantaciones, pero la pregunta acerca del impacto que tienen las plantaciones certificadas en sobre la conservación de la biodiversidad en regiones donde los bosques se han perdido por completo sigue sin respuesta. En este estudio, comparamos las comunidades de aves, mariposas y plantas en una región con alta deforestación en Santander, Colombia, con el fin de determinar el potencial para la conservación de dos tipos de certificaciones al café: Rainforest alliance y Rainforest alliance + Orgánico. Utilizamos 13 fincas, donde censamos aves, mariposas y plantas y además cuantificamos las características de la estructura del sombrío. No encontramos diferencias significativas en la mayoría de los índices de diversidad y composición de aves, mariposas y plantas entre diferentes tipos de cafetales. Sin embargo, a pesar de la alta variación entre las fincas, encontramos que la riqueza y abundancia de las mariposas aumentan con la disminución en el uso de pesticidas en las plantaciones. Estos resultados sugieren que el uso reducido de compuestos químicos en plantaciones certificadas promueven la conservación de las comunidades de mariposas. La biodiversidad asociada a estas plantaciones de café y las altas tasas de deforestación en Santander, sugieren que independientemente del tipo de certificación, estas plantaciones proveen los últimos refugios para la conservación de la biodiversidad en esta región.

Tropical Andes is one of the major biodiversity hotspots in which almost 7% of the total world’s endemic plants and 6% of terrestrial vertebrates are found (Myers *et al*. 2000). However, high deforestation rates are threatening the subsistence of this considerable portion of the world’s biodiversity (Rengifo *et al*. 1997, Brooks *et al*. 2002). The increase of urban areas and fast conversion of land for agricultural use are likely to be responsible for the transformation of the natural vegetation cover (Allen & Barnes 1985, Etter *et al*. 2006, Grau & Aide 2008). Coffee plantations may be a great contributor to the deforestation rate given that they occupy 44% of the total permanent crops in northern Latin America (Perfecto *et al*. 1996). These facts advocate for the development of sustainable alternatives to mitigate the detrimental effect caused by such practices and alleviate the extinction rates of local biodiversity (Rengifo *et al*. 1997, Brooks *et al*. 2002).

Shade coffee plantations might be a sustainable alternative because they support greater species richness of multiple groups of organisms than coffee grown in full sun (Perfecto *et al*. 1996, Wunderle Jr & Latta 1996, Greenberg *et al*. 1997a, Perfecto *et al*. 2007). In the past decade, research has focused on the differences in community composition and diversity between different types of shade coffee plantations, which are thought to contribute differentially to biodiversity conservation (Philpott *et al*. 2008). Birds, butterflies, ants and plants, are more diverse under less intensified shade coffee than mono-dominant shade plantations and in smaller households (Greenberg *et al*. 1997b, Mas & Dietsch 2003, Perfecto *et al*. 2003, Armbrecht *et al*. 2005, Philpott *et al*. 2007, Philpott *et al*. 2008, Mendez *et al*. 2010, Goodall *et al*. 2014). These studies suggest that the increased diversity is the result of differences in shade and socio-economical characteristics between coffee plantations, and some may therefore have a high potential value for biodiversity conservation (Philpott *et al*. 2008, Mendez *et al*. 2010, Goodall *et al*. 2014).

Lately, certification programs for shade coffee plantations have been implemented in several coffee-growing countries (Perfecto *et al*. 2005). Such certifications apply several standards for producing coffee in an attempt to make the process environmentally, socially and economically friendly (Perfecto *et al*. 2005, Philpott *et al*. 2007, Giovanucci *et al*. 2008, Philpott *et al*. 2008). Rainforest Alliance certified coffee, for example, has to be produced under a shade with foliage cover of at least 40%, represented by more than 80 trees and 12 native species per hectare and a double-layered canopy (Rice 2008, Sustainable Agricultural Network 2011). Also, Rainforest Alliance certified coffee-producing farms have to conserve forest remnants, reduce the use of chemical products, protect stream sides and encourage their workers to preserve the environment by proactive behavior (Rice 2008, Sustainable Agricultural Network 2011). Other types of certification programs such as organic certifications are not generally related to shade production, but prohibit the use of chemical products such as pesticides and/or fertilizers (Vandermeer 1995).

Although, these certification programs are presumably environmentally friendly, few empirical data have been gathered to test the hypothesis that increased diversity and complexity in shade cover increases biological diversity within coffee plantations (Hole *et al*. 2005, Philpott *et al*. 2008). Mas and Dietsch (2003, 2004) showed that bird and butterfly communities were more diverse in certified plantations that had a higher shade complexity (i.e. Bird Friendly Certification), but in another study, Philpott et al (2007) showed that diversity of birds and ants did not differ among plantations regardless of their shade complexity and whether they where certified or not. Both of these studies agree with the fact that no type of plantation sustains the diversity of natural forests, questioning the real usefulness of certification programs in preserving biodiversity (Mas & Dietsch 2003, O’Brien & Kinnaird 2003, Rappole *et al*. 2003, Mas & Dietsch 2004, Philpott *et al*. 2007). More recently, Blackman and Rivera (2011) showed that there is little evidence to support the claim that certification programs are beneficial for the environment. Given the disagreement among these results, additional studies are needed before general conclusions can be drawn, especially from regions of high conservation priority such as Colombia (Myers *et al*. 2000, Blackman & Rivera 2011). Additionally, all of these studies have focused on the levels of biodiversity sustained by certified coffee plantations only in comparison to natural forest remnants. Thus the question remains how might certification programs help preserve biodiversity in regions where natural forests have already been lost?

The Colombian Andes is a region with one of the highest conversion rates of natural ecosystems due to agricultural and demographic land use (Etter & van Wyngaarden 2000). In central Santander most of the landscape has been transformed to shade coffee plantations leaving few to no remnants of natural forests (Guhl 2009). In the past decade many farms have switched from highly intensified management of coffee to organic or certified programs, which implement standards to make coffee production environmentally friendly (Guhl 2009). This is one of Colombia´s oldest coffee-growing regions (Palacios 1980) and most of the coffee is grown under diverse and complex shade, making Santander Colombia´s greatest producer of certified coffee (Rainforest Alliance 2007, Guhl 2009, NaturaCert 2011). In other cases, farms may implement different types of shade such as shaded monoculture (as described by Moguel & Toledo 1999). These different types of shade programs found in the same geographic region allow a comparison of the role of different certification programs in biodiversity conservation.

In this study we test if farms with Rainforest Alliance and Rainforest Alliance plus organic certifications support a higher diversity of birds, butterflies, and trees than uncertified farms in the coffee-growing region of Santander. Specifically, we predicted that communities of such organisms inhabiting certified farms should be more diverse than those on non-certified farms. Additionally, shade structure and complexity should be higher in certified plantations and less complex in non-certified plantations.

## METHODS

### Study area and site selection

The coffee-growing region of Santander is located on the northwestern slope of the Eastern Andes in Colombia at middle elevations between 1200 and 1800 m. It is characterized by having a bimodal precipitation regime with dry seasons between July and August and December and January, with a mean annual precipitation of 1700 mm and a mean annual temperature of 20°C. Plantations in Santander differ from the rest of Colombia because almost all coffee is grown under a diverse shade that is typical of commercial polycultures (for detailed description of a commercial polyculture see Moguel and Toledo 1999). This type of plantation has high shade tree diversity but is dominated principally by legumes (*Inga* sp., *Erythrina* sp. and *Albizia* sp.), but some other non-legumes (*Cupania* sp., *Trichanthera gigantea*, *Cedrela* sp., *Tabebuia* sp.) are also relatively abundant. For decades farmers have kept this agricultural practice because the dry climatic conditions and high solar radiation otherwise make it impossible to grow coffee (Instituto Geográfico Agustin Codazzi 1996, Duran *et al*. 2004, Guhl 2009). For these reasons the region pioneered the production of certified coffee in Colombia and during the last decade Santander has became the most important producer of Rainforest Alliance certified coffee (Rainforest Alliance 2007, Guhl 2009, NaturaCert 2011). On the larger scale, coffee plantations dominate the landscape, along with cattle pastures and sugar cane, cassava and citrus plantations.

Within the municipalities of Páramo, Valle de San José, San Gil, and El Socorro, we selected 13 farms with at least 20 hectares of shaded coffee, except for two farms that had less planted area. The farms selected had to be either non-certified or certified by one of two types of certification: Rainforest Alliance certified (RA, hereafter) and Rainforest Alliance and organic certified (RA+Organic, hereafter) and had been certified for at least five years.

### ORGANISM AND SHADE STRUCTURE SAMPLING

Within each farm we identified up to ten points that were at least 100 meters apart from each other and 75 meters away from the plantation border, where we censed birds, tress and butterflies (for a detailed summary of sample sizes see Table 1). We carried out four censuses during two seasons: October - December 2005, March - May 2006. During each season, each point was visited twice for a 10-minute period. Censuses started at dawn and were conducted until 10:30 am. Every bird seen and heard within a 25-meter radius was counted, but flying and nocturnal birds were excluded from the analyses (Hutto *et al*. 1986). In each of the bird census points a 20 × 20 m vegetation plot was constructed to evaluate shade structure where we measured height, identity and diameter at breast height (DBH) of each tree as well as average canopy cover and tree density in each plot. Height was quantified using a clinometer, DBH was measured using a metric tape and canopy cover was recorded using a spherical densiometer. Finally, we sampled butterflies during the driest season of 2007 (July) and the wet season of 2008 (March) using standard Van Someren-Rydon traps in 24 points where birds and trees were censed. In each point we set up two traps, one placed in the canopy and the other one in the understory (above and below three meters) (DeVries 1987, 1988, DeVries *et al*. 1997). We used two types of baits, decomposing fish and fermenting bananas, in order to take into account different butterfly feeding guilds. These were assigned randomly to each trap in each point. We sampled each trap and bait was changed once every 48 hours for a period of eight days in each field season, for a total of eight samples per trap.

**Table 1.** Sample size and site selection used to evaluate the differences of bird, trees and butterfly communities between certified and non-certified coffee plantations.

|  | Certification | Number of Points Censused |  |  |
| --- | --- | --- | --- | --- |
|  |  | Birds | Trees | Butterflies |
| *San Francisco* | non-certified | 8 | 8 | 4 |
| *San Jose* | non-certified | 5 | 5 | 4 |
| *Damasco* | non-certified | 9 | 9 | N/A |
| *Montebello* | non-certified | 10 | 10 | N/A |
| *Total non-certified* |  | 32 | 32 | 8 |
| *San Fernando* | Rainforest Alliance | 10 | 10 | N/A |
| *Morros* | Rainforest Alliance | 10 | 10 | 4 |
| *La Pradera* | Rainforest Alliance | 9 | 9 | N/A |
| *Los Sinchos* | Rainforest Alliance | 10 | 10 | N/A |
| *La Laguna* | Rainforest Alliance | 10 | N/A | 4 |
| *Total Rainforest Alliance* |  | 49 | 39 | 8 |
| *Buenavista* | Rainforest + Organic | 10 | 10 | N/A |
| *El Galapo* | Rainforest + Organic | 9 | 9 | 4 |
| *El Placer* | Rainforest + Organic | 10 | 10 | N/A |
| *Transvaal* | Rainforest + Organic | 10 | 10 | 4 |
| *Total Rainforest+Organic* |  | 39 | 39 | 8 |
| *Total* |  | 110 | 110 | 24 |

### STATISTICAL ANALYSES

To compare species richness among coffee plantations we constructed abundance-based rarefaction curves using 100 bootstrap replicates (Gotelli & Colwell 2001). To statistically test for differences in species richness we rarefied the number of species to lowest number of individuals encountered across types of plantation. Additionally, we calculated Chao 1 species richness estimator, especially for butterfly richness where rarefaction curves did not appear to approach an asymptote. We computed the effective number of species based on the Shannon and Simpson indices, which take into account species richness and abundance. The effective number of species gives the number of species that a locality would have if all of the species would be represented by the same number of individuals. For the Shannon index, the effective number of species *x* is *x* = *e*^*shannon*^, and for the Gini-simpson index the effective number of species is 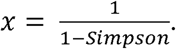 The correction to the indices allows us to directly interpret magnitude of the difference across types of plantations (Jost 2006). Finally we calculated similarity between farms using the Chao-Jaccard index, which accounts for unseen species (Chao *et al*. 2005). Using the Chao-Jaccard dissimilarity index we performed a Non-Metric Multidimensional Scaling (NMDS) to explore differences across farms. Diversity analyses were conducted the vegan package (Oksanen *et al*. 2013) for R (R Core Team 2013). We then compared if farms with a particular coffee type where more similar than farms across coffee types using a nonparametric ANOVA (Kruskall Wallis test) and an Analysis of similarity (ANOSIM). To test for differences in the shade structure (DBH, tree density, canopy height, canopy cover and management index) among types of plantations, we used one-way ANOVAs and post-hoc Tukey tests for those that showed significant differences after testing for the normality of the variables.

Using the vegan package, we fitted five species abundance distributions models to the data for each type of plantation: Null, Geometric series, Lognormal, Zipf and Mandelbrot. The models differ in the shape of the curve mainly in predicting the number of species with intermediate and low abundances. The fitting of such models allows us to evaluate the for the Gini-simpson index the effective number of species is similarity in community composition based on abundance and dominance patterns. We selected the model that fits our data the best by using AIC.

We used a published database to classify bird species in twelve diet guilds that represent the preferences of species (Karr *et al*. 1990). Such guilds were: FRI (Frugivores insectivores), F (Frugivores), CF (Canopy Frugivores), UF (Understory Frugivores), GR, (Granivores), I (Insectivores), CI (Canopy Insectivores), UI (Understory Insectivores), NEC (Nectarivores), NI (Nectarivores Insectivores), OM (Omnivores), RAP (Raptors). We performed the guild analysis only for birds because there are no published datasets for the guilds of adult butterflies. Statistical significance was assessed by estimating the 95% confidence interval for the mean; if confidence intervals overlapped then, we assumed no differences in the mean number of species or individuals in each guild.

Following Mas and Dietsch (2003), we calculated a Management Index for each vegetation plot sampled using four variables: mean diameter at breast height (DBH), number of tree species tree density and mean tree height. This Management Index quantifies the level of intensification that each certification has over the shade canopy and it is based on the values that each variable measured has in a native forest. Because we did not have a native forest for comparison, we assumed the maximum value of each variable in the entire sample plots to indicate low intensification and recall “natural conditions”. Thus, by comparing the least intensified value of each variable to the observed value in each plot we calculated the amount of management a plot has relative to the expected “natural conditions”. For each variable, the index varies between 0.0 and 1.0, where 0.0 means the plot is similar in that variable to presumed “natural conditions” and 1.0 means that the plot is highly intensified. After calculating the amount of management for each variable, all of these values were added to obtain the Management Index of the plot. We only included four variables in the index: DBH, number of tree species, tree density and tree height, because we did not have canopy cover values for all of the plots. The Management Index for the plot therefore potentially varied between 0.0 and 4.0. We then used the mean values of the Management Index to compare the intensification between the types of plantations studied using a one-way ANOVA. Further, to evaluate if the level of intensification in the farm had an effect on birds and butterflies we calculated linear regressions between bird and butterfly richness, and abundance in each farm and the Management Index.

## RESULTS

RA and RA + Organic farms had more butterfly and tree species than Non-Certified farms but differences where not significant (Kruskal-Wallis: Butterflies: Chi-squared = 4.57, df = 2, p = 0.1; Trees: Chi-squared = 1.53, df = 2, p = 0.46; Table 2; Figure 1; Table S1). In contrast, Non-Certified coffee plantations had more bird species than do certified plantations but again differences where not significant (Kruskall-Wallis: Chi-squared = 0.35, df = 2, p = 0.8; Table 2; Figure 1). There was no significant difference in the estimated or rarefied richness of birds, butterflies or trees across types of plantations (Kruskall-Wallis: Estimated richness: Birds: Chi-squared = 1.1, df = 2, p = 0.6; Butterflies: Chi-squared = 3.7, df = 2, p = 0.15; Trees: Chi-squared = 2.9, df = 2, p = 0.23; Rarefied Richness: Birds: Chi-squared = 1.01, df = 2, p = 0.6; Butterflies: Chi-squared = 0.85, df = 2, p = 0.65; Trees: Chi-squared = 1.52, df = 2, p = 0.46). Again, there was no difference in the effective number of species across types of plantations (Kruskall-Wallis: Shannon: Birds: Chi-squared = 1.2, df = 2, p = 0.6; Butterflies: Chi-squared = 3.42, df = 2, p = 0.18; Trees: Chi-squared = 1.95, df = 2, p = 0.37; Simpson: Birds: Chi-squared = 2.1, df = 2, p = 0.35; Butterflies: Chi-squared = 2, df = 2, p = 0.36; Trees: Chi-squared = 2.4, df = 2, p = 0.3). However, the overall abundance of butterflies was five times higher in RA + Organic plantations than in Non-Certified plantations and almost twice as high as RA plantations. When lumping farms across types of plantations, there was no difference in the rarefied number of bird species but RA and RA + Organic resulted to be more species rich in trees and butterflies than non-certified plantations (Figure 1; Statistical significance is given by the non-overlap in the 95% confidence intervals).

**FIGURE 1.**
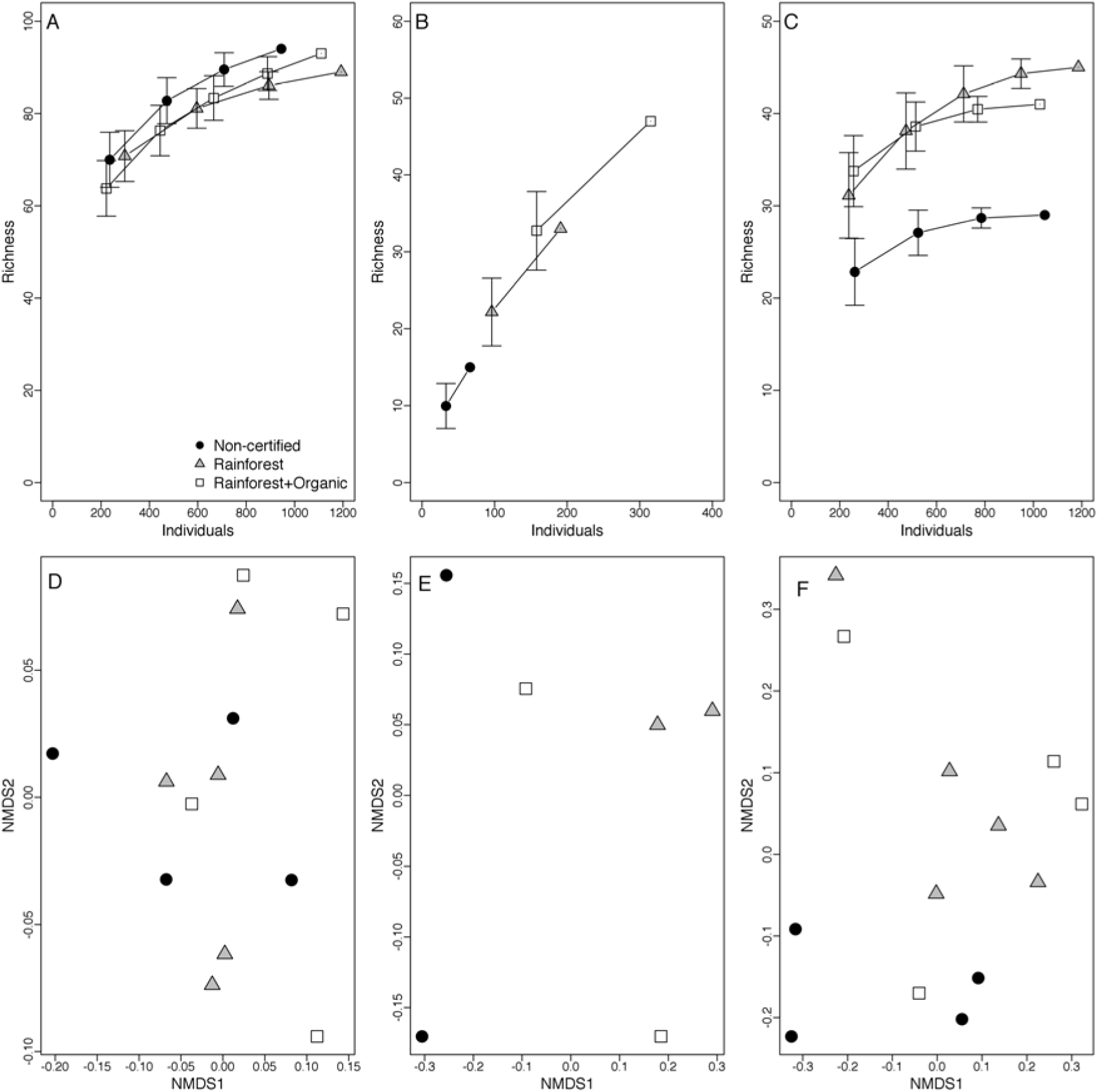
Species accumulation curves (upper panels) and NMDS of A,D) birds, B,E) butterflies and C,F) trees in Rainforest+Organic, Rainforest and non-certified plantations in Santander. Error bars represent the 95% confidence intervals of the rarefied estimated number of species.

We found no significant difference in community structure between plantations. Chao-Jaccard index did not show a higher species turnover between plantations across types (Kruskal-Wallis; Birds: Chi-squared = 1.4, df = 1, p = 0.23; Butterflies: Chi-squared = 0.18, df = 1, p = 0.66; Trees: Chi-squared = 0.45, df = 1, p = 0.49). This was corroborated by the ANOSIM. The only difference is that RA + Organic certified farms resulted to be significantly different among them in bird composition (ANOSIM: Bids; R = -0.17, p = 0.04; Butterflies: R = 0.16, p = 0.52; Trees: R = 0.09, p = 0.21). Bird community structure resulted to be similar according to the models fitted to the rank-abundance distributions. For all three types of plantations, the best model was the Preemption model with exactly the same slope (Figure 2A; Table S2). In turn, Non-Certified butterfly and plant communities resulted to follow a Zipf model and RA and RA + Organic butterfly communities followed a Mandelbrot and Zipf models. The best model that fitted plant communities in both types of certified plantations was a lognormal model (Figure 2; Table S2). Finally, we found no difference in the bird guild structure of birds across types of plantations. The distributions of number of species and proportional number of individuals were almost identical among types of plantations (Figure 3; Statistical significance given by the 95% confidence interval of the mean shown by the error bars). There are some differences between species richness and abundance of the guilds across plantations. Even though Insectivorous birds were more species rich; Frugivorous and Insectivorous birds were more abundant (Figure 3). We provide complete species lists for birds, trees, and butterflies present in each type of plantation in Table S3, S4 and S5.

**FIGURE 2.**
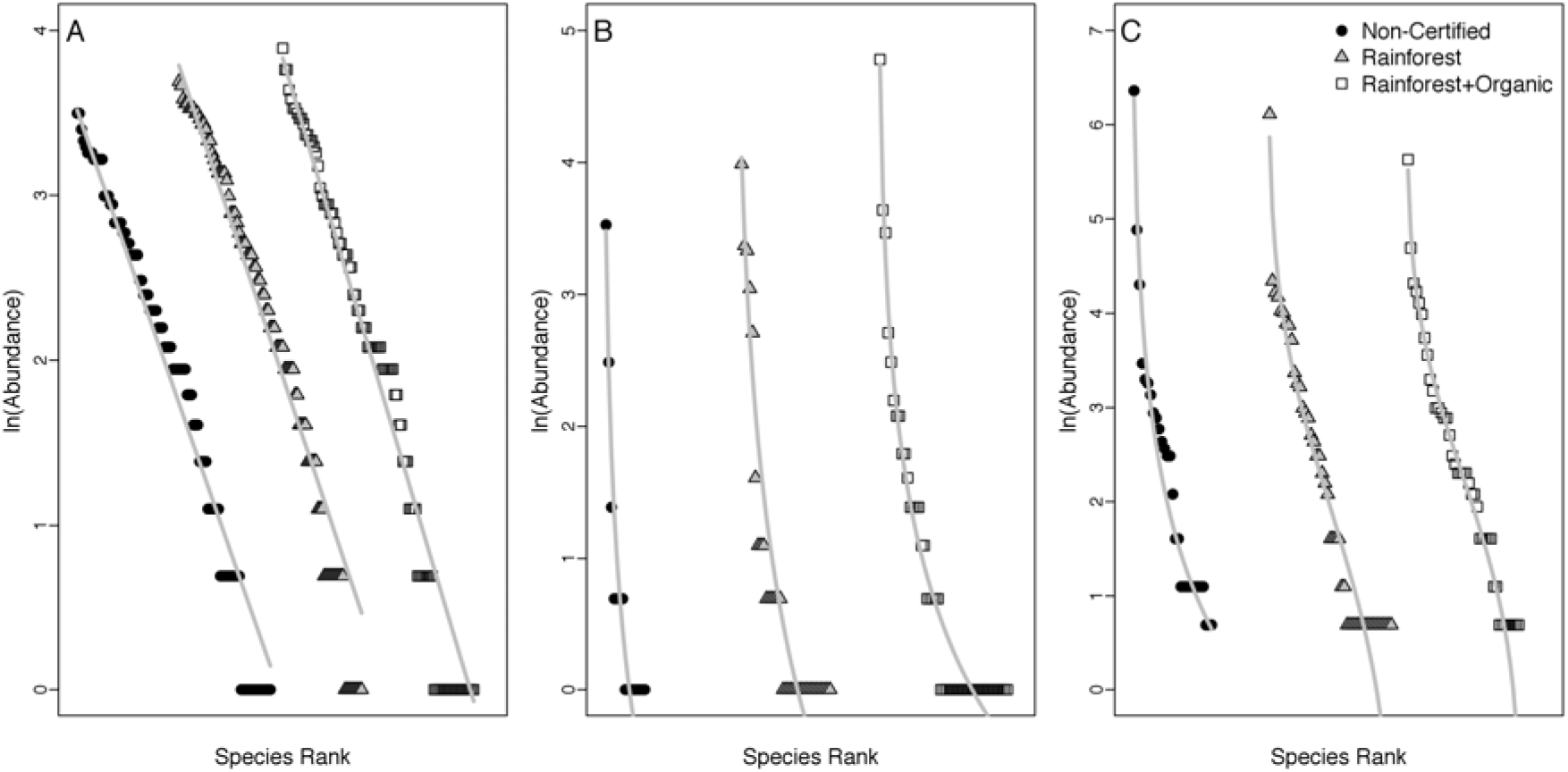
Species rank curves for A) Birds, B) Butterflies and C) Trees in Non-certified, Rainforest and Rainforest+Organic certified plantations. The gray line shows the predictions of the best fit model (Full results of the models are shown on Table S2)

**FIGURE 3.**
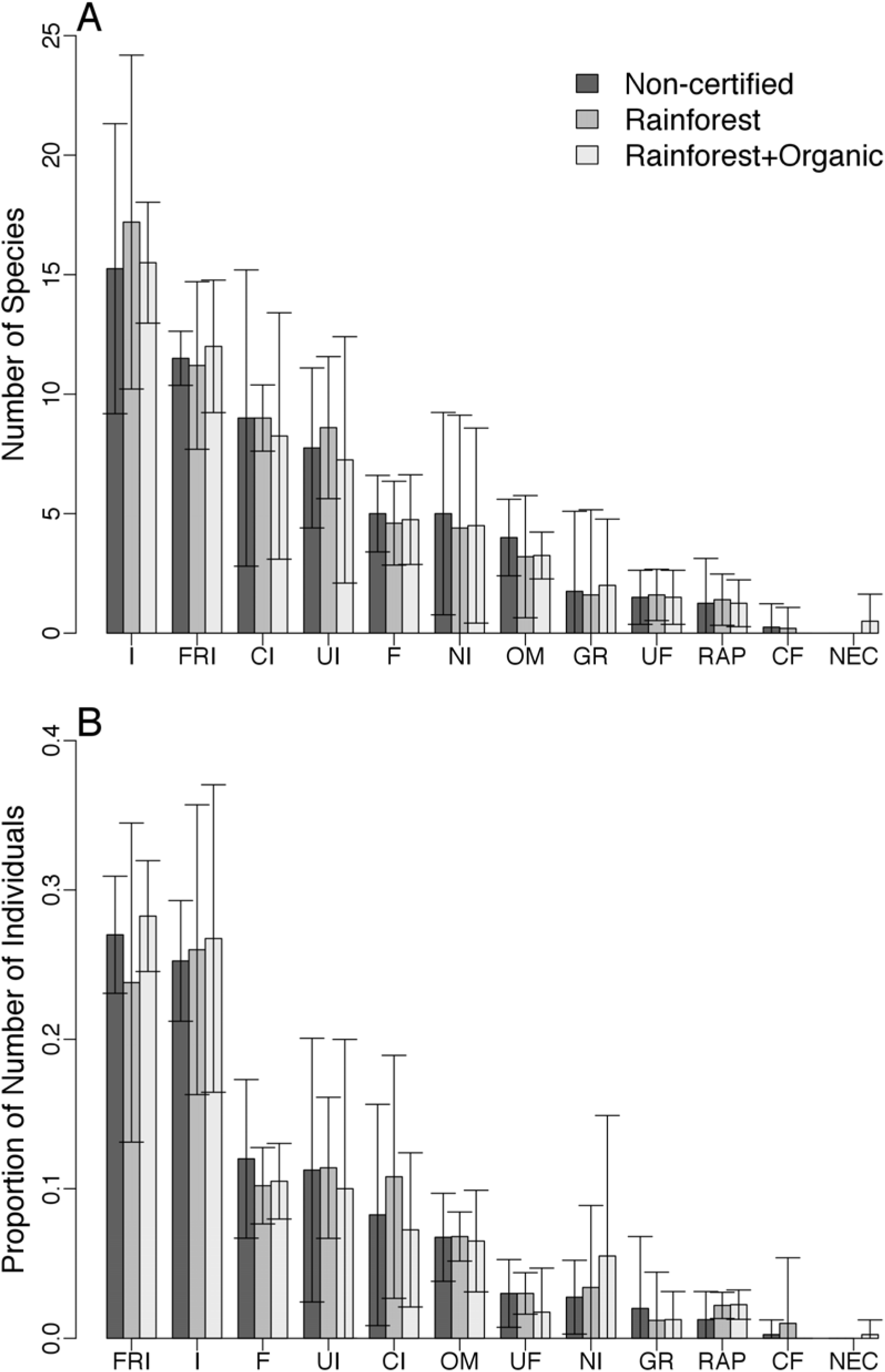
A) Average number of species and B) average proportion of individuals in each diet guild for the three types of coffee plantations. Error bars indicate 95% confidence intervals of the mean. I (Insectivores), FRI (Frugivores insectivores), CI (Canopy Insectivores), UI (Understory Insectivores), F (Frugivores), NI (Nectarivores Insectivores), OM (Omnivores), GR (Granivores), UF (Understory Frugivores), RAP (Raptors), CF (Canopy Frugivores), NEC (Nectarivores).

Shade structure was similar between the three types of coffee plantations (Table 3). Specifically, we found no significant differences in DBH (ANOVA; n = 13, f = 0.67, p = 0.52), canopy cover (ANOVA; n = 12, f = 0.72, p = 0.51), tree density (ANOVA; n = 13, f = 1.66, p = 0.23) or tree height (ANOVA; n =13, f = 3.69 p = 0.06). When summarizing all variables into the Management Index, we found that RA+Organic had the lowest and the non-certified plantations the highest values. Generally a high value of the Management Index is associated with a decrease in the shade cover. However, the differences between certified and non-certified plantations where not statistically significant (ANOVA; n = 13, F = 2.63 p = 0.11). Finally, butterfly and bird species richness and abundance were not related to management index (Table S6).

## DISCUSSION

Overall, we found no evidence that RA and organic certification are better refuges for biodiversity than non-certified farms. Instead, certified and non-certified plantations provide similar types of habitat that can serve as refuge to biodiversity in a region where almost all of the forest has been lost. Even though we found no significant differences in birds, butterfly and tree richness or abundance it is interesting to note the steep increase in butterfly overall richness. This increase suggests that the high variance in Butterfly diversity between coffee plantations might drive the absence of statistical significance (See Table S1). The results are in line with previous studies that have reported that butterfly abundance and richness decrease sharply with increasing plantation intensification and less intensified plantations tend to have mores species rich plant communities (Botero & Baker 2001, Mas & Dietsch 2003, Perfecto *et al*. 2003, Mas & Dietsch 2004). Birds showed an opposite trend to our predictions, suggesting that factors other than shade structure and agrochemical use dominate the response of birds to these types of conservation actions. Potentially, an increase in bird species diversity might depend on the landscape matrix composition and the amount of forest available around the plantations (Sisk *et al*. 1997). Alternatively, this result might suggest that some certified plantations do not have the complexity necessary to improve bird conservation (Philpott *et al*. 2007).

Overall we were surprised that most of our comparisons resulted to be non-significant, especially in butterfly estimated richness and diversity. The sampling effort was exactly the same for all of the farms and even then, we found a five fold increase in the number of butterflies captures between Non-certified and RA + Organic farms. Great variance and low sample size might have contributed on the absence of significant differences in abundance and richness. The absence in turnover patters is most likely the result of butterflies in Non-certified plantations are just a subset of the species from the other types of plantations that can overcome pesticides or are more generalists in their larval stages. Thus, no turnover is observed but just addition of new species with practices promoted by certification programs.

### Overall Community Structure

The fact that bird communities follow a Preemption model suggests that there is a high dominance of one or few species and the rest of the species are represented by few individuals (Motomura 1932; Figure 2; Table S2). However, the effective number of species suggests that the dominance is not very extreme given by the differences between observed and effective species number (Table 2). In contrast, models of butterfly communities suggest that for all communities, there are niche mechanisms operating in the abundance of species. The Zipf and Mandelbrot models assume that some species have a priori niche requirements for their presence and establishment in the communities (Frontier 1985, Wilson 1991). Pioneer species have little requirements in the community while species from later successional stages need higher complexity in niche and resource availability. Thus, late successional species should be represented by a lower number of individuals within the community (Frontier 1985, Wilson 1991). Such result suggests that butterfly communities in coffee plantations may be remnants of the diversity lost by deforestation even in Non-Certified plantations. However, effective species number suggests a great inequality in the abundance of species. Such high inequality is corroborated by the low values of the third parameter in the Mandelbrot model of the RA plantations (Wilson 1991). The results are intriguing and further research comparing forest and plantation butterfly diversity might clarify our results. Unfortunately, the total absence of forest in the region did not allow us to comply with this.

**Table 2.** Mean ± 1 standard deviation in *abundance, richness and diversity of bird, butterfly and tree communities in non-certified and certified coffee plantations. No. of species and No. of individuals correspond to the observed data and Chao corresponds to the estimated number of species, Rarefied # of Species corresponds to the number of species rarefied to the smallest number observed in a plantation and Exp (Shannon) and 1/(1-Simpson) correspond to the effective species number based on the Shannon and Simpson diversity indices*.

|  | No.<br>Species | No.<br>Individuals | Chao | Rarefied # Sp. | Exp (Shannon) | 1/(1-Simpson) |
| --- | --- | --- | --- | --- | --- | --- |
| **Birds** |  |  |  |  |  |  |
| Non-Certified | 62 ± 8 | 236 ± 73 | 75 ± 12 | 53 ± 3 | 51.2 ± 6 | 45.02 ± 4 |
| Rainforest | 63 ± 10 | 263 ± 83 | 79 ± 14 | 51 ± 4 | 49.4 ± 9 | 42.43 ± 9 |
| Rainforest+Organic | 61 ± 6 | 246 ± 42 | 71 ± 9 | 50 ± 2 | 47.44 ± 4 | 40.43 ± 4 |
| **Tree** |  |  |  |  |  |  |
| Non-Certified | 12 ± 6 | 262 ± 80 | 12 ± 8 | 11 ± 5 | 5 ± 3 | 3.2 ± 2 |
| Rainforest | 18 ± 6 | 237 ± 47 | 18 ± 8 | 17 ± 7 | 10 ± 5 | 6.0 ± 3 |
| Rainforest+Organic | 18 ± 7 | 256 ± 88 | 18 ± 11 | 17 ± 6 | 11 ± 6 | 7.9 ± 9.5 |
| **Butterflies** |  |  |  |  |  |  |
| Non-Certified | 10 ± 1 | 33 ± 16 | 15 ± 3 | 8 ± 2 | 4.6 ± 1 | 2.96 ± 1 |
| Rainforest | 21 ± 2 | 96 ± 16 | 34 ± 4 | 9 ± 2 | 10.1 ± 3 | 6.87 ± 2 |
| Rainforest+Organic | 29 ± 8 | 158 ± 29 | 52 ± 28 | 10 ± 3 | 11.7 ± 8 | 6.93 ± 6 |

Plant communities in certified plantations follow a lognormal distribution (Figure 2; Table S2). The lognormal distribution is characterized by low dominance and high number of species with intermediate abundance. Most likely this is the product of the shade plantation process and a certification effect in which shade species are selected to have about the same abundance per hectare to comply with certification requirements. Few species such as *Inga sp*. have high abundance in these plantations because they are planted in a slightly higher density to improve nitrogen fixation and soil productivity.

### Bcird guild structure

The fact that bird guild structure was identical across types of plantations suggests that the slight differences in shade structure have little influence on bird diversity. An interesting fact is a decoupling in the richness and abundance patterns. Types of plantations have slightly different order of most abundant groups. For example in Non-certified RA + Organic plantations the most abundant group was the Frugivorous Insectivorous and in the RA the most abundant was the Insectivorous. However, the small differences in abundance between types of plantations suggest that the mechanisms driving the abundance of species is homogeneous across plantations. The fact that the insectivorous birds dominate the community might me explained by the dominance of *Inga sp*. trees in the canopy. These trees have extrafloral nectaries that attract high abundance and diversity of insects.

### Relationship between butterfly and plant communities

Our results regarding the differences in Management Index are consistent with the hypothesis and previous evidence that differences in shade structure between plantations should drive the increase in butterfly diversity (Mas & Dietsch 2004). Despite that most of the results in butterfly diversity do not significant differences, richness, abundance and composition suggest the opposite (Figure 1, Figure 2, Table 2, Table S2). Other studies have reported a relationship between shade characteristics and the diversity of organisms in coffee plantations and our results conform to the pattern of decreased butterfly diversity in more intensified plantations (Greenberg *et al*. 1997b, Soto-Pinto *et al*. 2001, Perfecto *et al*. 2003, Armbrecht *et al*. 2005, Philpott *et al*. 2008). However, it is unlikely that differences in shade structure alone cause such an increase in the diversity of butterfly communities, especially if the differences between the types of plantations where not significant. It is probable that differences in tree species richness also play an important role in explaining these differences. We found that there was a higher number of tree species in RA+Organic than those in non-certified plantations (Table 2). Thus, because of strong dependency of butterflies on their host plants (Ehrlich & Raven 1964), absolute plant species richness might explain the differences in butterfly community composition between coffee plantations. Additionally, the preservation of streamsides and herbaceous vegetation (as required by RA certification) could enhance the diversity of butterflies by protecting their host plants in the certified plantations and creating suitable habitats for these insects (Sparks & Parish 1995).

An alternative explanation for the differences in butterfly communities between the types of plantations is the decreased use of pesticides and chemical fertilizers in certified coffee plantations. Pesticides, herbicides and chemical fertilizers might directly affect butterflies by killing the adults or immature stages, or indirectly, by reducing the availability of host plants (Longley & Sotherton 1997). In fact, RA certified farms, which allow moderate use of such chemical compounds, were intermediate in diversity of butterflies. Consequently, our results might also be consistent with the latter explanation; nevertheless a combination of the two factors could result in the increase of butterfly diversity in certified plantations (Weibull *et al*. 2000). This result provides evidence to the positive effect of the certification programs on butterfly diversity conservation, and to potential benefits to the coffee growers in terms of enhanced profits through improved ecosystem services provided by butterflies (Tscharntke *et al*. 2005, Sandhu *et al*. 2008). For example, high butterfly diversity might directly benefit the plantations as pollination or weed control agents and indirectly as potential food resources for birds that might enhance biological control of insect pests or seed dispersal (Perfecto *et al*. 2004, Tscharntke *et al*. 2005, Klein *et al*. 2007, Sandhu *et al*. 2008).

### Tree communities and shade structure

Shade structure as interpreted by the Management Index between plantations was nearly significantly different between RA+Organic and non-certified plantations. However, this pattern is more likely to be driven by the differences in species richness and height of the shade. The rest of the variables were similar between plantations suggesting that the structure of the shade is equivalent among treatments (Table 3). The similarity in shade structure might be the result of climate in the localities sampled. These localities have the highest levels of light radiation in Colombia and have low levels of rainfall and overall humidity (Instituto Geográfico Agustin Codazzi 1996). These environmental conditions force coffee growers to maintain higher tree density and canopy cover in coffee plantations even if they are not certified (Duran *et al*. 2004, Guhl 2009). Non-certified plantations had decreased diversity of shade tree species and a higher abundance of *Inga* sp. trees. *Inga* spp. trees provide a high quality of nectar in their flowers and extra floral nectaries that attract a high number of insects, and insectivorous and nectarivorous birds (Calvo & Blake 1998, Johnson 2000). This might explain why we found higher bird species richness in non-certified plantations, although the differences between plantations were not significant.

**Table 3.** Management index and shade structure values of each type of plantation.

|  | Management | Canopy |  | Canopy |  |
| --- | --- | --- | --- | --- | --- |
|  | Index | Height (m) | DBH (cm) | Cover (%) | Tree Density |
| *Non-Certified* | $2.85 \pm 0.92$ | $10.51 \pm 4.6$ | $22.86 \pm 7.23$ | $83.45 \pm 10.22$ | $10.32 \pm 5.64$ |
| *Rainforest* | $2.44 \pm 0.42$ | $13.44 \pm 1.36$ | $24.76 \pm 5.15$ | $89.00 \pm 4.18$ | $10.96 \pm 3.43$ |
| *Rainforest+Organic* | $2.43 \pm 0.53$ | $12.45 \pm 2.1$ | $24.4 \pm 5.28$ | $84.87 \pm 3.89$ | $11.97 \pm 2.72$ |

### Implications for conservation

This region of Santander has been highly affected by deforestation, resulting in a landscape dominated by pastures, coffee and sugar cane plantations. Such conversions occurred during the beginning of the 1900s leaving the region with no natural forests for a long period of time (Federación Nacional de Cafeteros 1957, Palacios 1980). This loss is observed in the absence of common forest bird species such as woodcreepers (Furnariidae) and manakins (Pipridae) that are present in this type of agroecosystem elsewhere (Greenberg *et al*. 1997a, Greenberg *et al*. 1997b, Calvo & Blake 1998, Johnson 2000, Perfecto *et al*. 2003). Despite the evidence that vegetation factors influence the diversity of birds in coffee plantations we were unable to find this relationship in our study sites (Philpott *et al*. 2008). This might also reflect the loss of source populations of many forest species that might benefit from environmentally friendly practices of coffee certification. As an evidence for this, we found that most of the bird species inhabiting the three types of plantations have low sensitivity to human impact and are able to nest in or prefer secondary vegetation habitats (Figure S1). However, the high diversity of migrant species (21 species of forest nearctic migrants), the presence of several species that prefer forest habitats and have high sensitivity to human impact, and the presence of endemic and highly threatened species such as Niceforo’s Wren, Chestnut-bellied Hummingbird, Golden-winged Warbler and Cerulean Warbler, increase the value of these coffee plantations and motivate coffee growers to improve the habitats for these species (Table S3).

Mid-elevations in the northern Andes have been subject to high deforestation rates probably because of their productive soils and favorable climate (Myers *et al*. 2000). In the Colombian Andes, more than 70% of the natural cover has been transformed to land for agricultural use (Cavelier 1997). One of the main uses of this agricultural land is coffee plantations which represent about 34% of the total permanent crop area, 6% of the total export products and contributes almost 1.8% of Colombia’s gross domestic product (Espinal *et al*. 2005). Despite the fact that coffee production in Colombia has decreased in the last decade, the total area occupied by plantations has increased (Figure S2). These statistics suggest that it is highly unlikely that conversion of natural forests to coffee plantations will stop or even decrease. In this case, shade coffee plantations might be the only alternative to preserve the native biodiversity that still persists in these regions. In fact, Santander is the only region in Colombia in which shade coffee is growing faster than pasture or any other type of crops (N. Galindo *pers. comm*.).

In some cases, farmers obtain certification status with little changes to their crops. Climatic conditions in the area force the coffee growers to maintain a dense and diverse shade that is further co-opted to fulfill the requirements of the RA certification. However, both organic and RA certification programs radically change the management of the plantation by reducing and prohibiting the use of chemical fertilizers and pesticides. Other features that are radically changed from the farms managements are the protection of stream-sides and remnant natural vegetation with in the farm. It is very likely that without the certification programs, such fragments and water protection is lost to coffee bushes for increased farm production. In this way, promoting the improvement of shade coffee plantations will enhance the conservation of biodiversity that is already happening in Santander.

While we chose the farms to be homogeneous in size for appropriate comparison of the levels of biodiversity sustained by each type of plantation, the region is characterized by a high variance in farm size and socio-economical status of the landowners. However, as a program of the Colombian Coffee Federation (FNC), the landowners receive assistance for certification as groups of producers irrespective of the size of the land. Thus, the certification is not applied only to the farm but to a group of nearby farms that are randomly inspected for certification. Recent studies suggest that smaller coffee plantations might preserve higher levels of biodiversity (Mendez *et al*. 2010, Goodall *et al*. 2014). In the case of this particular region such fact might not apply because of the way FNC provide support to the growers. However, a formal analysis to evaluate the influence of farm size on biodiversity conservation in this region is still needed. Unfortunately, we do not have the appropriate information at hand to provide some insight about the size influence. However, we might speculate that the model of cooperative certification as used in the farms evaluated by the present study might increase the impact of bigger farms to preserve biodiversity conservation (Mendez *et al*. 2010, Goodall *et al*. 2014) by homogenizing practices across producers in the same group.

Because of the uniqueness of coffee production in Santander, our results have to be carefully interpreted. The climatic conditions and low soil productivity, compared to other regions in Colombia, forces coffee growers to create deeper topsoil that enhances coffee production. This means that, overall, shade-grown coffee plantations in Santander are managed less intensively than in other places in Colombia or Latin America. These facts perhaps explain why the impact of the certification programs on biodiversity was less than might have been expected.

## Acknowledgements

We would like to thank Fundación Natura, Cenicafé and the Tropical Andean Butterfly Diversity Project for financing this study. The Coffee Growers Committee of Santander, Henry Parra, Nelson Galindo and Fernando López for their help during the planning of the study and field seasons. Luis Quintero and Paula Sarmiento for their assistance in the field. The coffee growers that allowed us to work in their farms; the Santos family and Jorge Julian Santos for their hospitality and collaboration during this work. S. K. Robinson, J. Ungvari-Martin, K. R. Willmott, P. C. Zalamea, P.R. Stevenson and two anonymous reviewers for their valuable comments on the final manuscript.

## References

Allen, J. C., and D. F. Barnes. 1985. The Causes of Deforestation in Developing Countries. Annals of the Association of American Geographers 75: 163-184.

Armbrecht, I., L. Rivera, and I. Perfecto. 2005. Reduced diversity and complexity in the leaf-litter ant assemblage of Colombian coffee plantations. Conservation Biology 19: 897-907.

Blackman, A., and J. Rivera. 2011. Producer-Level Benefits of Sustainability Certification. Conservation Biology 25: 1176-1185.

Botero, J. E., and P. S. Baker. 2001. Coffee and biodiversity: a producer-country perspective. *In* P. S. Baker (Ed.). Coffee futures: a source book of some critical issues confronting the coffee industry, pp. 94-103. CAB international, Wallingford, UK.

Brooks, T. M., R. A. Mittermeier, C. G. Mittermeier, G. A. B. Da Fonseca, A. B. Rylands, W. R. Konstant, P. Flick, J. Pilgrim, S. Oldfield, G. Magin, and C. Hiltontaylor. 2002. Habitat Loss and Extinction in the Hotspots of Biodiversity Pérdida de Hábitat y Extinciones en Áreas Críticas para la Biodiversidad. Conservation Biology 16: 909-923.

Calvo, L., and J. Blake. 1998. Bird diversity and abundance on two different shade coffee plantations in Guatemala. Bird Conservation International 8: 297-308.

Cavelier, J. 1997. Selvas y Bosque montanos. *In* M. E. Chaves and N. Arango (Eds.). Informe nacional sobre el estado de la biodiversidad. Instituto de Investigación de Recursos Biológicos Alexander von Humboldt, Bogotá.

Chao, A., R. L. Chazdon, R. K. Colwell, and T. J. Shen. 2005. A new statistical approach for assessing similarity of species composition with incidence and abundance data. Ecology Letters 8: 148-159.

Devries, P. J. 1987. The Butterflies of Costa Rica and Their Natural History Volume I: Papilionidae, Pieridae, Nymphalidae. Princeton University Press, Princeton.

Devries, P. J. 1988. Stratification of fruit-feeding nymphalid butterflies in a Costa Rican rainforest. Journal of Research on the Lepidoptera 26: 98-108.

Devries, P. J., D. Murray, and R. Lande. 1997. Species diversity in vertical, horizontal, and temporal dimensions of a fruit-feeding butterfly community in an Ecuadorian rainforest. Biological Journal of the Linnean Society 62.

Duran, S. M., R. Garcia, J. G. Velez, O. A. Echeverry, and J. E. Botero. 2004. Caracterización de la biodiversidad en paisajes rurales cafeteros. Informe técnico preliminar. Ventana No. 3: Santander. p. 114. Cenicafé, Chinchiná.

Ehrlich, P. R., and P. H. Raven. 1964. Butterflies and Plants: a study in coevolution. Evolution 18: 586-608.

Espinal, C. F., H. J. Martinez, and X. Acevedo. 2005. La cadena del Café de Colombia: una mirada global de su estructura dinámica 1991 - 2005. Ministerio de Agricultura y desarrollo Territorial.

Etter, A., C. McAlpine, K. Wilson, S. Phinn, and H. Possingham. 2006. Regional patterns of agricultural land use and deforestation in Colombia. Agriculture, Ecosystems & Environment 114: 369-386.

Etter, A., and W. Van Wyngaarden. 2000. Patterns of landscape transformation in Colombia, with emphasis in the andean region. AMBIO: A Journal of the Human Environment 29: 432-439.

Federación Nacional De Cafeteros. 1957. Origen del cultivo del café en los varios departamentos del país. Agricultura Tropical 13: 326-334.

Frontier, S. 1985. Diversity and Structure in Aquatic Ecosystems. Oceanogr Mar Biol 23: 253-312.

Giovanucci, D., P. Liu, and A. Byers. 2008. Adding Value: certified coffee trade in North America. *In* P. Liu (Ed.). Value-adding Standards in the North American Food market - trade opportunites in certified products for developing countries. FAO, Rome.

Goodall, K. E., C. M. Bacon, and V. E. Mendez 2014. Shade tree diversity, carbon sequestration, and epiphyte presence in coffee agroecosystems: A decade of smallholder management in San Ramón, Nicaragua. Agriculture, Ecosystems and Environment 199: 200-206.

Gotelli, N. J., and R. K. Colwell. 2001. Quantifying biodiversity: procedures and pitfalls in the measurement and comparison of species richness. Ecology Letters 4: 379-391.

Grau, H. R., and M. Aide. 2008. Globalization and Land-Use Transitions in Latin America.

Greenberg, R., P. Bichier, A. C. Angon, and R. Reitsma. 1997a. Bird populations in shade and sun coffee plantations in central Guatemala. Conservation Biology 11: 448-459.

Greenberg, R., P. Bichier, and J. Sterling. 1997b. Bird populations in rustic and planted shade coffee plantations of eastern Chiapas, México. Biotropica 29: 501-514.

Guhl, A. 2009. Café bosques y certificación agricola en Aratoca, Santander. Revista de Estudios Sociales - Universidad de los Andes 32: 114-125.

Hole, D. G., A. J. Perkins, J. D. Wilson, I. H. Alexander, P. V. Grice, and A. D. Evans. 2005. Does organic farming benefit biodiversity? Biological Conservation 122: 113-130.

Hutto, R. L., S. M. Pletschet, and P. Hendricks. 1986. A fixed-radius point count method for nonbreeding and breeding season use. The Auk 103: 593-602.

Instituto Geográfico Agustin Codazzi. 1996. Diccionario geográfico de Colombia. *In* IGAC (Ed.). IGAC., Bogotá.

Johnson, M. D. 2000. Effects of shade-tree species and crop structure on the winter arthropod and bird communities in a Jamaican shade coffee Plantation. Biotropica 32: 133-145.

Jost, L. 2006. Entropy and diversity. Oikos 113: 363-375.

Karr, J. R., S. K. Robinson, J. G. Blake, and O. Bierregaard. 1990. Birds of four neotropical forests. *In* A. W. Gentry (Ed.). Four Neotropical Rainforests. Yale University Press, New Haven, Connetticut, USA.

Klein, A.-M., B. E. Vaissière, J. H. Cane, I. Steffan-Dewenter, S. A. Cunningham, C. Kremen, and T. Tscharntke. 2007. Importance of pollinators in changing landscapes for world crops. Proceedings of the Royal Society B: Biological Sciences 274: 303-313.

Longley, M., and N. W. Sotherton. 1997. Factors determining the effects of pesticides upon butterflies inhabiting arable farmland. Agriculture, Ecosystems & Environment 61: 1-12.

Mas, A. H., and T. V. Dietsch. 2003. An index of management intensity for coffee agroecosystems to evaluate butterfly species richness. Ecological Applications 13: 1491-1501.

Mas, A. H., and T. V. Dietsch. 2004. Linking shade coffee certification to biodiversity conservation: butterflies and birds in Chiapas, Mexico. Ecological Applications 14: 642-654.

Mendez, V. E., C. M. Bacon, M. Olson, K. S. Morris, and A. Shattuck. 2010. Agrobiodiversity and Shade Coffee Smallholder Livelihoods: A Review and Synthesis of Ten Years of Research in Central America. Prof Geogr 62: 357-376.

Moguel, P., and V. M. Toledo. 1999. Review: biodiversity conservation in traditional coffee systems of Mexico. Conservation Biology 13: 11-21.

Motomura, I. 1932. A statistical treatment of associations. Japanese Journal of Zoology 44: 379-383.

Myers, N., R. A. Mittermeier, C. G. Mittermeier, G. A. Da Fonseca, and J. Kent. 2000. Biodiversity hotspots for conservation priorities. Nature 403: 853-858.

Naturacert. 2011. Estadisticas de produccion de café por departamento.

O’Brien, T., and M. Kinnaird. 2003. Caffeine and conservation. Science 300: 587-587.

Oksanen, J., F. G. Blanchet, R. Kindt, P. Legendre, P. R. Minchin, R. B. O’HARA, G. L. Simpson, P. Solymos, M. H. Stevens, and H. Wagner. 2013. vegan: Community Ecology Package.

Palacios, M. 1980. Coffee in Colombia, 1850 - 1970. An economic, social an political history. Cambridge University Press, Cambirdge.

Perfecto, I., I. Armbrecht, S. M. Philpott, T. V. Diestch, and L. Soto-Pinto. 2007. Shaded coffee and the stability of rainforest margins in Latin America. *In* T. Tscharntke, C. Leuschner, M. Zeller, E. Guhudja and A. Bidin (Eds.). The stability of tropical rainforest margins: linking ecological, economic, and social constraints of land use and conservation Springer, Environmental Science Series, Heidelberg, Germany.

Perfecto, I., A. Mas, T. Dietsch, and J. Vandermeer. 2003. Conservation of biodiversity in coffee agroecosystems: a tri-taxa comparison in southern Mexico. Biodiversity and Conservation 12: 1239-1252.

Perfecto, I., R. A. Rice, R. Greenberg, and M. E. V. D. Voort. 1996. Shade coffee: a disappearing refuge for biodiversity. BioScience 46: 598-608.

Perfecto, I., J. Evandermeer, A. Mas, and L. Pinto. 2005. Biodiversity, yield, and shade coffee certification. Ecological Economics 54: 435-446.

Perfecto, I., J. H. Vandermeer, G. L. Bautista, G. I. Nuñez, R. Greenberg, P. Bichier, and S. Langridge. 2004. Greater predation in shaded coffee farms: the role of resident neotropical birds. Ecology 85: 2677-2681.

Philpott, S. M., W. J. Arendt, I. Armbrecht, P. Bichier, T. V. Diestch, C. Gordon, R. Greenberg, I. Perfecto, R. Reynoso-Santos, L. Soto-Pinto, C. Tejeda-Cruz, G. Williams-Linera, J. Valenzuela, and J. M. Zolotoff. 2008. Biodiversity loss in Latin American coffee landscapes: review of the evidence on Ants, Birds, and Trees. Conservation Biology 22: 1093-1105.

Philpott, S. M., P. Bichier, R. Rice, and Rainforest ALLIANCE. 2007. Field-testing ecological and economic benefits of coffee certification programs. Conservation Biology 21: 975-985.

R Core Team. 2013. R: A language and environment for statistical computing. R Foundation fro Statistical Computing, Vienna, Austria.

Rainforest Alliance. 2007. Summary of certified area under cultivation.

Rappole, J., King, and J. Vega-Rivera 2003. Coffee and conservation. Conservation Biology 17: 3.

Rengifo, L. M., G. P. Servat, J. M. Goerck, B. A. Loiselle, and J. G. Blake. 1997. Patterns of Species Composition and Endemism in the Northern Neotropics: A Case for Conservation of Montane Avifaunas. Ornithological Monograpphs 48 Studies in Neotropical Ornithology Honoring Ted Parker: 577-594.

Rice, R. 2008. Drinking green. A primer on choosing coffee that supports sustainable practices. Smithsonian Zoogoer July - August: 17 - 23.

Sandhu, H. S., S. D. Wratten, R. Cullen, and B. Case. 2008. The future of farming: the value of ecosystem services in conventional and organic arable land. An experimental approach. Ecological Economics 64: 835-848.

Sisk, T. D., N. M. Haddad, and P. R. Ehrlich. 1997. Bird Assemblages in Patchy Woodlands: Modeling the Effects of Edge and Matrix Habitats. Ecological Applications 7: 1170-1180.

Soto-Pinto, L., Y. Romero-Alvarado, J. Caballero-Nieto, and G. Segura Warnholtz. 2001. Woody plan diversity and structure of shade-grown-coffee plantations in Northern Chiapas, Mexico. Revista de Biología Tropical 49: 977-987.

Sparks, T. H., and T. Parish. 1995. Factors affecting the abundance of butterflies in field boundaries in Swavesey fens, Cambridgeshire, UK. Biological Conservation 73: 221-227.

Sustainable Agricultural Network. 2011. Group Certification Standard. http://www.sanstandards.org.

Tscharntke., T., A. M. Klein., A. Kruess., I. Steffan-Dewenter., and C. Thies. 2005. Landscape perspectives on agricultural intensification and biodiversity – ecosystem service management. Ecology Letters 8: 857-874.

Vandermeer., J. 1995. The Ecological basis of alternative agriculture. Annual Review of Ecology and Systematics 26: 201-224.

Weibull., A.-C., J. Bengtsson., and E. Nohlgren. 2000. Diversity of butterflies in the agricultural landscape: the role of farming system and landscape heterogeneity. Ecography 23: 743-750.

Wilson., J. B. 1991. Methods for Fitting Dominance Diversity Curves. J Veg Sci 2: 35-46.

Wunderle Jr., J., and S. Latta. 1996. Avian abundance in sun and shade coffee plantations and remnant pine forest in the Cordillera Central, Dominican Republic. Ornitologia Neotropical 7: 19-34.

